# microRNA-721 is a host regulator of TNF-IRF1 axis in *Leishmania* infected macrophage

**DOI:** 10.64898/2026.05.13.724987

**Authors:** Jonathan Miguel Zanatta, Ian Antunes Ferreira Bahia, Emanuella Sarmento Alho de Sousa, Clara Andrade Teixeira, Kathia Terumi Kato, Camilla de Almeida Bento, Stephanie Maia Acuña, Maria Regina D’Império Lima, Ricardo Silvestre, Dennyson Leandro M Fonseca, Sandra Marcia Muxel

**Affiliations:** Laboratory of Molecular Immunology Regulation (miRLab), School of Arts, Science and Humanities (EACH), University of São Paulo (USP), São Paulo, SP, Brazil; Postgraduate Program on Immunology, Department of Immunology, Institute of Biomedical Sciences (ICB), University of Sao Paulo (USP), São Paulo, SP, Brazil; Life and Health Sciences Research Institute (ICVS), School of Medicine, University of Minho, Braga, Portugal; ICVS/3B’s - PT Government Associate Laboratory, Braga/Guimarães, Portugal

**Keywords:** Macrophage, Transcriptome, microRNAs, Leishmaniasis

## Abstract

MicroRNAs (miRNAs) are small noncoding RNAs that play critical roles in regulating immune responses and have emerged as potential biomarkers and therapeutic targets in complex diseases. Leishmaniasis is a neglected disease that compromises host immunity and is associated with challenging treatments regimens. *Leishmania amazonensis* (*L. amazonensis*), an intracellular protozoan parasite, causes cutaneous leishmaniasis by replicating inside mammalian macrophages to establish infection. In this context, miRNAs have emerged as vital post-transcriptional factors that regulate the inflammatory landscape during infection.

In this study, we aimed to analyze the function of miR-721 in macrophages during *L. amazonensis* infection by integrating in silico miR-721 target prediction with RNAseq data from macrophages of two distinct mouse genotypes, resistant C57BL/6 and susceptible BALB/c. We found that miR-721 is induced in macrophages infected with *L. amazonensis*, but is not in LPS-stimulated macrophages, suggesting a TLR4-independent activation. Integrating miR-721 target prediction with comparative transcriptomic analyses in resistant C57BL/6 and susceptible BALB/c models revealed the TNF-IRF1 axis as a primary miR-721-associated regulatory network. Specifically, miR-721 is predicted to target the 3’UTRs of *Tnf* and *Irf1* to suppress the inflammatory response. Functional inhibition of miR-721 successfully restored *Tnf* and *Irf1* expression and reduced the amastigote burden over 24 hours. Furthermore, we showed that the miR-721/TNF-IRF1 axis regulates downstream genes associated with macrophage response, such as *Serpine1*, *Csf1*, *Cd*69 and *Maf*. Our work demonstrated that *Leishmania* induces miR-721, which negatively modulates the TNF-IRF1 axis, thereby suppressing the immune response and favoring parasite persistence. While C57BL/6 macrophages exhibit a robust activation of the TNF-IRF1 network, promoting inflammatory response, BALB/c macrophage showed a breakdown of this network. This was associated with post-transcriptional suppression of inflammatory responses, thereby favoring parasite persistence. These findings link miR-721 to the establishment of macrophage polarization, providing relevant insights into the mechanisms of parasite subversion of the host immune response.

## 1 Introduction

Leishmaniasis is a parasitic disease that affects more than one million people worldwide each year and remains a major public health problem in endemic regions^1,2^. Cutaneous leishmaniasis is a clinical manifestation of infection caused by *Leishmania amazonensis*, particularly in the Amazon region of Brazil^1,2^. In cutaneous leishmaniasis, macrophages are central to disease pathogenesis because they act both as effector cells and as the major intracellular niche for *Leishmania* survival and replication^3–6^. Upon parasite recognition, macrophages activate inflammatory pathways that promote phagocytosis, cytokine production, and microbicidal activity^3–6^. However, once internalized, *Leishmania* can subvert macrophage activation, impair antigen presentation^7,8^, and attenuate secondary stimulation^8,9^, resulting in a restrained inflammatory response thereby favoring parasite persistence^7,10,11^.

The outcome of *Leishmania* infection is therefore linked to how efficiently macrophages sustain inflammatory and antimicrobial transcriptional programs^12^. Among the molecules involved in this context, tumor necrosis factor (TNF) plays a central role in macrophage activation and leishmanicidal activity^13^. The transcription factor interferon regulatory factor 1 (IRF1) is also induced and drives the expression of genes involved in antimicrobial activity, including pathways related to nitric oxide (NO) and TNF production^14,15^. Thus, suppression of TNF and IRF1-mediated responses may represent an important mechanism by which *Leishmania* establishes infection and persists within host cells^16^.

In addition to transcriptional regulation, macrophage responses to *Leishmania* are also shaped by microRNAs (miRNAs), which regulate gene expression at the post-transcriptional level and have emerged as important modulators of host-pathogen interactions^17,18^. Previous studies, including ours, have shown that miR-721 is induced in macrophages during *Leishmania* infection and modulates nitric oxide biosynthesis^19–23^. However, the functional role of miR-721 in *Leishmania*-infected macrophages remains poorly understood.

Here, we investigated the contribution of miR-721 to macrophage transcriptional responses during *L. amazonensis* infection. By integrating miR-721 target prediction with comparative transcriptomic analyses of infected macrophages from resistant (C57BL/6) and susceptible (BALB/c) mouse strains, we identified miR-721-associated genes centered on *Tnf* and *Irf1* expression. Functional analysis highlighted the role of miR-721 in the regulation of the TNF-IRF1 axis in macrophages during *L. amazonensis* infection.

## 2 Methods

### 2.1 Ethics statement

All experiments using animals were conducted in accordance with ethical guidelines for the care and use of laboratory animals established by São Paulo state (Law11.977/2008) and Brazilian Federal Government (Law 11.794/2008). The study protocol was approved by the Institutional Animal Care and Use Committee (CEUA) of the Institute of Biomedical Science (ICB) from University of São Paulo (approval number CEUA-ICB: CEUA N° 5090100822). For bone marrow cell isolation, mice were euthanized via isoflurane inhalation, strictly adhering to institutional safety and welfare policies.

### 2.2 RNA-seq and differential expression genes

RNA-seq data were obtained from Sequence Read Archive (SRA), under accession PRJNA481042, comprising *Mus musculus* Bone Marrow Derived Macrophages (BMDMs) from C57BL/6 and BALB/c mice infected with *L. amazonensis* (24 hours post infection) or corresponding uninfected controls^19,24^. Differential expression analyses were performed for each mouse strain (C57BL/6 and BALB/c) using the R DESeq2 package^25,26^. In this context, Differentially Expressed Genes (DEGs) were defined using False Discovery Rate (FDR) threshold < 0.01, and Log 2-Fold Change (log2FC) values were considered < 0 as down-regulated and > 0 as up-regulated. To visualize all results, we used the ggplot2 R package^27^. In addition, all subsequently analysis was performed in R programing language^28^.

### 2.2 Identification and validation of miR-721 targets

To characterize the miR-721-associated regulatory landscape, we employed an integrative approach combining target prediction with differential expression analysis. Candidate targets were retrieved using the R package multiMir^29^, aggregating predictions from multiple databases. To increase the robustness of predicted findings, only validated genes or those with high prediction score (> 0.7) were retained, prioritizing high-confidence interactions and reducing noise from low-probability predictions^29^. In this context, the resulting predicted microRNAs Targetome was subsequently intersect with experimental RNA-seq data form *L. amazonensis*-infected C57BL/6 and BALB/c macrophages.

### 2.3 Rank-based feature selection

To evaluate the ability of miR-721–associated genes to discriminate infected and control macrophages, we first performed principal component analysis (PCA) using the filtered gene sets. We then prioritized genes according to the magnitude of differential expression, ranking them by absolute log2 fold change (|Log2FC|)^25,30^. To emphasize gene with stronger expression changes, we calculated a weighted contribution based squared Log2FC values and retained the subset of genes accounting for 50% of the cumulative contribution. This approach allowed us to reduce the gene set while prioritizing the most strongly modulated features for downstream analyses^31^. Gene sets from each strain were subsequently combined for downstream analyses.

### 2.4 Enrichment analysis

To investigate the biological functions associated with selected miR-721 targets, we conducted a functional enrichment analysis using the clusterProfiler R package and org.Mm.eg.db to mouse annotations^32–35^. This analysis allowed us to identify enriched genes sets associated with biological processes (BPs) ontology. Statistical significance for enriched terms was set at an FDR < 0.05.

### 2.5 Protein-protein interaction networks

Protein–protein interaction (PPI) analysis was constructed using the STRING database via the STRINGdb R package (version 12)^36^, restricted to *Mus musculus* (taxonomy ID: 10090), with a confidence score threshold of 400 (medium confidence)^37^. Interaction networks were generated using DEGs from the complete transcriptomic dataset without target-based filtering. Networks were visualized to highlight interactions involving key genes, and Circos plots were generated to represent the interaction landscape.

### 2.6 Parasite culture

*L. amazonensis* promastigotes (strain MHOM/BR/1973/M2269) were maintained in M199 medium (Gibco, ThermoScientific, USA) supplemented with 10% heat-inactivated fetal bovine serum (Invitrogen), 5mg/L hemine, 100 μM adenine, 100 U penicillin, 100 μg/mL streptomycin, 40 mM Hepes-NaOH, and 12 mM NaHCO_3_ (SigmaAldrich, USA). Culture was incubated at 25°C and mxaintained at low passage numbers (up to passage 5) to preserve virulence^19,38–41^. Growth cycles were monitored weekly to ensure the parasites reached the stationary-phase for experimental infection.

### 2.7 Macrophage cell culture and differentiation

Infection and stimulation assays were performed using either the RAW 264.7 macrophages cell line (ATCC) or primary BMDMs. For primary cultures, bone marrow cells were isolated from femurs and tibiae of 6-to-8-week-old female BALB/c and C57BL/6 mice^19,38–41^. These animals were supplied and maintained by the Animal Facility Center of the Institute of Biomedical Science, USP. Macrophage was differentiated using bone marrow-derived cells cultured in L929-conditioned medium (10%) per 7 days^19,38–41^. Cells were maintained in RPMI 1640 medium (LGC, Brazil), supplemented with 10% inactivated fetal bovine serum, 50 U/mL penicillin, 50 μg/mL streptomycin (Gibco, Thermo Scientific, USA),) at 37° C and 5% CO_2_^19,38–41^.

### 2.8 Infection quantification

After macrophage differentiation, cells were seeded into 8-well chamber slides (5×10^4^ cells/well; Millipore, Merck, Darmstadt, Germany). For the infection analysis, *L. amazonensis* promastigotes in the stationary growth phase were added to the cultures at parasite-to-host (MOI) of 5:1. Following 4 h of infection, cultures were washed twice with 1X PBS pH 7.2, to remove non-phagocytosed parasites. After 4 and 24 h of infection, the chamber slides were washed two times with PBS 1X and were then fixed with acetone/methanol (1:1 *v/v;* Sigma-Aldrich, St. Louis, MO, USA) and were incubated at -20 °C for 24 h followed by Giemsa staining (Fast Panotic Kit, Laborclin, São Paulo, Brazil). At least 600 macrophages were counted per well to achieve the parameters of the Macrophage Infection Rate [(infected macrophage/total macrophage) * 100], amastigotes per infected macrophage, and the infection index, which was calculated by multiplying the rate of infected macrophages by the average number of amastigotes per macrophage^19,38–41^.

### 2.9 *In vitro* infection and LPS stimulation for RT-qPCR analysis

Macrophages were seeded at density 3×10^6^ cells/well in 6-well plates and incubated overnight at 37°C in a 5% CO_2_ to allow for cell adherence. For the infection assay, *L. amazonensis* promastigotes in the stationary growth phase were added to the cultures at parasite-to-host (MOI) of 5:1. Following 4 h of infection, cultures were washed twice with 1X PBS pH 7.2, to remove non-phagocytosed parasites. Infected cells were then harvested or maintained in fresh RPMI medium for up to 24 h.

Macrophages were treated with 100 ng/mL of *Escherichia coli* LPS (serotype 0127: B8; Sigma-Aldrich, St. Louis, MO) by replacing the culture supernatant with LPS-supplemented RPMI medium for 4 h and 24 h, as described above. Non-infected and non-stimulated macrophages were maintained in culture in the control periods of the experiments.

### 2.10 Total RNA extraction

After infection or LPS stimulation, macrophages were collected in TRIzol Reagent® (Ambiom, ThermoScientific, USA) with the addition of 1X PBS, in the proportion of 750µL:250µL, respectively. RNA extraction was performed with the miRNeasy kit (QIAGEN, USA), following the manufacturer’s instructions. The concentration and quality of the RNA obtained were analyzed using spectrophotometry (Nanodrop).

### 2.11 Reverse Transcription Reaction and Real-Time PCR(RT-qPCR) for mRNA

The cDNA of mRNAs was obtained by reverse transcription reaction with 2 µg of total RNA and RevertAid Reverse Transcriptase enzyme and Random Hexamer Primer (Thermo Scientific, USA), to a final volume of 40 μL, following the cDNA synthesis protocol provided by the manufacturer’s instructions. Real-time RT-qPCR was performed on a StepONE Thermal Cycler (Thermo Scientific, USA). For the reaction, we used 10 μL of SYBR Green PCR Master Mix (Thermo Scientific, USA), 0.2 μM of oligonucleotide primers (Thermo Scientific, USA), 5 μL of template cDNA (diluted 100X) and water for a final volume of 20 μL/well. The reactions comprise the first cycle of 50°C for 2 min and the second cycle of 94°C for 10 min, followed by 40 cycles of 94°C for 30 s and 60°C for 30s. The primer list was as follows: *B2M*-F: 5′-cactgaattcacccccactga-3′ and *B2M*-R: 5′-acagatggagcgtccagaaag-3′; *Tnf*-F: 5′-ccaccacgctcttctgtcta-3′ and *Tnf*-R: 5′-agggtctgggccatagaact-3′; *Irf1*-F: 5′-caaccaaatcccagggctga-3′ and *Irf1*-R: 5′-cggaacagacaggcatcctt-3′; *Cux1*-F: 5′-atgaccgaccttgaacgagc-3′ and *Cux1*-R: 5′-ctggagccttctggatctgc-3′; and *Nos2*-F: 5′-agagccacagtcctctttgc-3′ and *Nos2*-R: 5′-gctcctcttccaaggtgctt-3′. The fold-change was calculated using the Delta-Delta Ct (ΔΔCt) method and was based on β-2-microglobulin was normalized with the uninfected group or negative control. The fold-change was presented in log2.

### 2.12 Reverse Transcription Reaction and Real-Time PCR (RT-qPCR) for microRNA

For miRNA analysis, cDNA was synthesized from 250 ng of total RNA using the miScript II RT Kit (Qiagen, Hilden, Germany) according to the manufacturer’s instructions. The reverse transcription reaction was performed in a final volume of 10 μL and included 5X miScript HiSpec Buffer, 10X Nucleics Mix, miScript Reverse Transcriptase Mix, and RNase-free water. The reaction was incubated at 37°C for 60 min, followed by enzyme inactivation at 95°C for 5 min. RT-qPCR was performed on a StepOne Thermal Cycler (Thermo Scientific, USA). For the reaction, we used 10 μL of miScript SYBR Green (Qiagen), 10X miScript Universal Primer, 10X specific primer, 2 μL of cDNA (10X diluted) and RNase-free water to a final volume of 10 μL/well. The reactions comprise the first cycle of 95°C for 15 s followed 40 cycles of 94°C for 15 s, 55°C for 30 s and 70°C for 30 s. Relative expression was normalized to SNORD95 and calculated using the ΔΔCt method. The fold-change was presented in log2.

### 2.13 miR721-mRNA interactions in *Tnf* and *Irf1*

*In silico* prediction of miR-721 target sites was performed using the miRmap software^42^, which integrates multiple robust prediction features, such as target site accessibility, conservation and score metrics, to more solid estimate and evaluate miRNA-mediated direct binding probability.

### 2.14 Inhibition of miRNA

Macrophages (3 x 10^6^) were plated in 6-well-plate and, after 24 h, transfected with miRVana miR-721 inhibitor or a Negative Control (NC) (ThermoScientific, USA) at a final concentration of 30 nM, combined with 3 µL of FuGENE (Promega, USA) in 200 µL of RPMI medium with 10% FBS, and previously incubated for 20 min at room temperature. Afterwards, the mix was transferred to cells washed twice with RPMI, incubated for 24 h and infected as described above.

### 2.15 Statistical analysis

Statistical analyses were performed according to data distribution, assessed by the Shapiro–Wilk test and quantile–quantile (QQ) plot inspection, with normality assumed for *p* > 0.05 and when data were within the QQ confidence intervals. For normally distributed data, comparisons between two groups were performed using Student’s t-test, while comparisons among two or more groups were conducted using one-way Analysis of Variance (ANOVA) followed by Sidak’s post hoc test. Effect sizes were estimated using Cohen’s d (**Supplementary Table S1**; **Supplementary Figures 1** and **2**). All analyses were primarily performed using the stats package^43^ in R (version 4.4.3), and statistical significance was defined as *p*-value < 0.05.

## 3 Results

### 3.1 miR-721 regulatory signatures in macrophages identifies *Tnf* and *Irf1* as key associated genes

To identify high-confidence regulatory targets of miR-721 during *L. amazonensis* infection, we used an integrative approach combining *in silico* miR-721-target prediction and transcriptomic profiling (**Figure 1a**). An initial target analysis identified 2,135 putative miR-721 target genes (**Supplementary Table S2**; **Supplementary Figure 3**). These predicted targets were subsequently integrated with RNA-seq data generated from BMDMs from C57BL/6 and BALB/c mice after 24 h of infection *in vitro* with *L. amazonensis*. Overall, differential expression analysis showed a total of 4.987 and 1.200 DEGs for C57BL/6 and BALB/c, respectively. In this context, from total DEGs for C57BL/6 and BALB/c (**Supplementary Table S3**; **Supplementary Figure 4)** the data were filtered to retain only genes predicted as miR-721 targets. Accordingly, C57BL/6 macrophages displayed 302 miR-721–associated DEGs, including 157 up and 145 down regulated, whereas BALB/c macrophages showed fewer target DEGs, with 28 up and 22 down regulated genes (**Figure 1b**).

**Figure 1.**
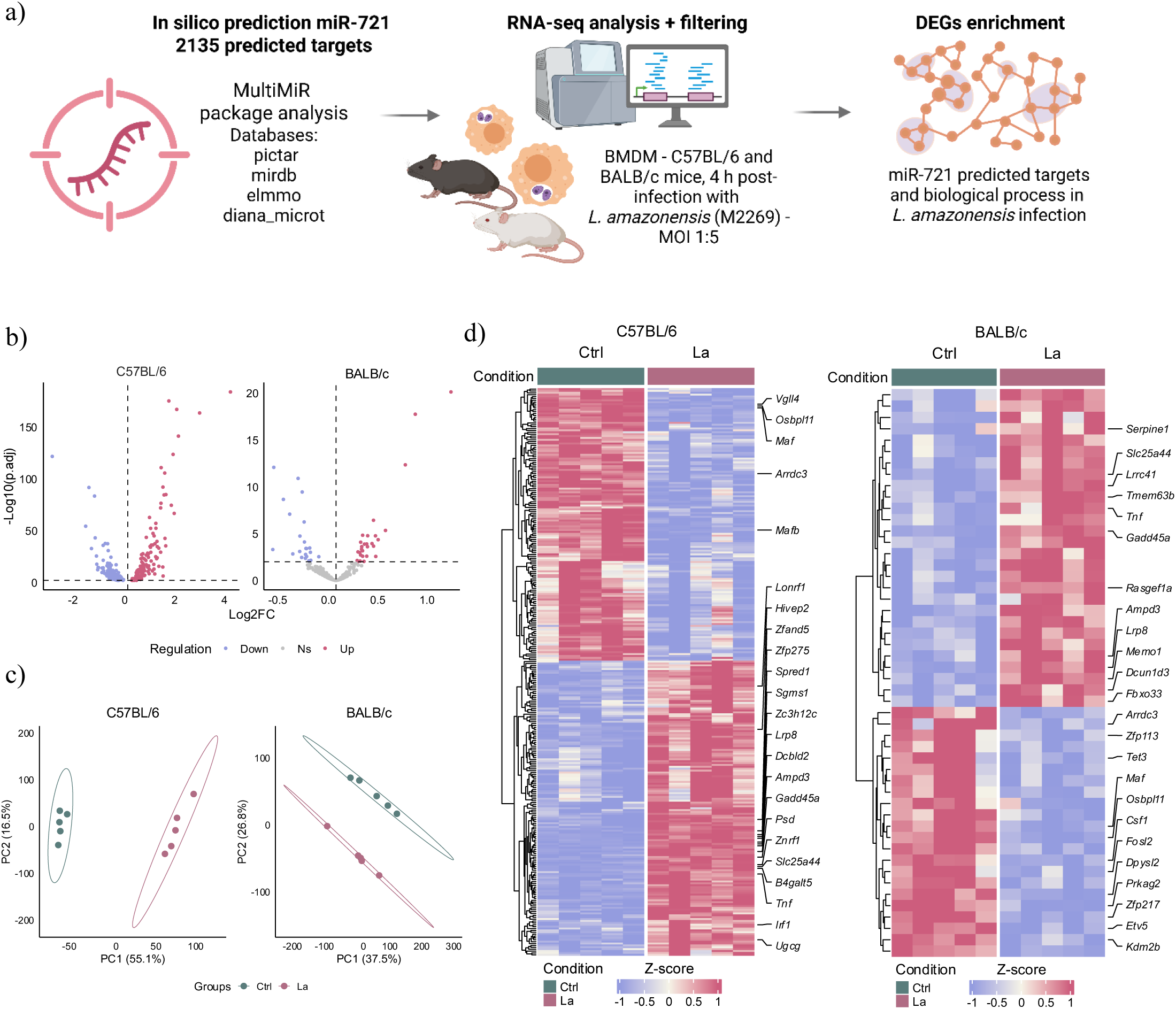
Integrative analysis identifies miR-721-associated differentially expressed genes in *L. amazonensis*-infected macrophages. **a)** Schematic overview of the analytical pipeline. **b)** volcanos plots showing the distribution of differentially expressed genes (DEGs) associated with predicted miR-721 from C57BL/6 and BALB/c mice. The red and blue colors represented up and down regulated genes respectively. **c**) Principal component analysis (PCA) of the filtered miR-721 associated DGEs sets showing the separation between control (green) and *L. amazonensis*-infected (pink) macrophages in each mouse strain. **d**) Heatmaps showing the expression patterns of miR-721-associated DEGs in control and infected macrophages from C57BL/6 and BALB/c mice. Rows represent genes and columns represent samples. Expression values are shown as Z-scores.

Furthermore, PCA was performed to evaluate the performance of miR-721–associated DEGs to discriminate between experimental groups. Variance-explained profiles and elbow plots showed that most of the variance in the miR-721-filtered expression space was captured by the first principal components (PCs). The results revealed that in C57BL/6 macrophages, the first two PCs together accounted for 90.1% of the total variance, with PC1 driving the clear separation between *L. amazonensis*–infected and control samples. A comparable separation pattern was observed in BALB/c macrophages, in which PC1 and PC2 together explained 75.9% of the variance and again distinguished infected conditions from control conditions (**Figure 1c; Supplementary Figure 5**). In this context, a rank-based feature selection analysis was conducted to define a more concise and informative set of genes (**Supplementary Tables S4-S6**), revealing shared DEGs between different mouse strains. Among these, *Tnf*, *Slc25a44*, *Arrdc3*, *Ampd3*, *Osbpl11*, *Maf*, *Lrp8*, and *Gadd45a* exhibited consistent expression patterns after infection in both C57BL/6 and BALB/c strains (**Figure 1d, Supplementary Table S7**).

To functionally characterize the signatures associated with the commonly shared miR-721 target DEGs, we performed a Gene Ontology (GO) enrichment analysis. The enriched biological processes (BP) were predominantly related to immune and inflammatory responses, including cytokine production regulation, innate immune signaling, and cellular response to tumor necrosis factor (**Figure 2a**; **Supplementary Tables S8**). Notably, genes such as *Tnf* and *Lrp8* exhibited similar up-regulated expression patterns in the C57BL/6 and BALB/c macrophages (**Figure 2b**). Furthermore, this BPs were predominantly enriched by *Tnf* and *Irf1* in C57BL/6 macrophages, whereas in BALB/c macrophages the enrichment was mainly centered on *Tnf*, highlighting a distinct organization of inflammatory networks associated with *L. amazonensis* infection (**Figure 2c, Supplementary Table S9**). Based on these observations, we hypothesized that miR-721 potentially regulates key targets such as *Tnf and Irf1*, thereby contributing to the modulation of the inflammatory responses during *L. amazonensis* infection.

**Figure 2.**
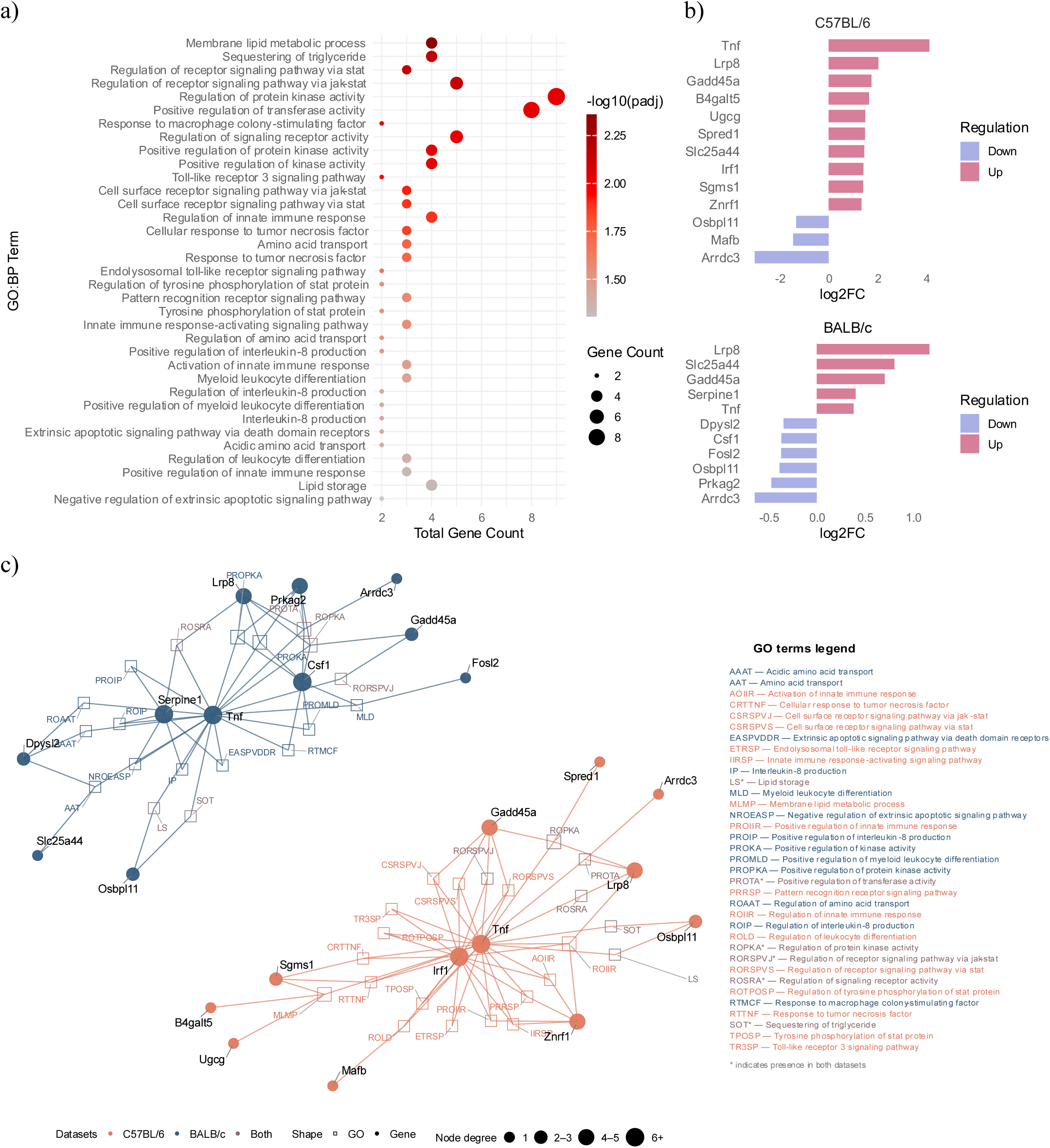
Functional enrichment analysis highlights pathways associated with miR-721-related genes during *L. amazonensis* infection. **a)** Dotplot shown enrichment analysis of shared miR-721-associated expressed genes. Dot size represent the gene count for each term, and color intensity indicates –log10 of adjusted *p-*value. **b)** Barplots showing enriched-gene expression levels for C57BL/6 and BALB/c. Upregulated genes are shown in pink, and down regulated genes are represented by blue. **c**) The network depicts the interaction between genes and enrichment sets. The circular nodes represent genes, and the square nodes represent pathways. Additionally, the orange and dark blue colors represent C57BL/6 and BALB/c, respectively. The size of the circular nodes indicates the degree of connectivity.

### 3.2 *L. amazonensis* infection induces miR-721 expression and correlates with modulation of inflammatory gene activation

To investigate the relationship between miR-721 and the inflammatory regulators identified in transcriptomic analysis, we quantified miR-721 and key genes expression in BMDMs from C57BL/6 and BALB/c mice under infection (La) and inflammatory (LPS) conditions at 4 h and 24 h timepoints (**Figure 3a**). Infection resulted in approximately 70% and 45% of infected macrophages, and amastigote-to-macrophage ratios ranging from 2.2 and 3-3.8 in C57BL/6 and BALB/c macrophages, respectively (**Figure 3b**). Following evaluation of infection parameters, RT-qPCR analyses revealed that miR-721 expression was significantly induced in both C57BL/6 and BALB/c macrophages during *L. amazonensis* infection. This increase was observed at both 4 h and 24 h post-infection when compared to uninfected control BMDMs (Ctrl) and LPS-treated groups (**Figure 3c-d**). To further explore miR-721 regulation, we assessed the expression of *Cut*-like homeobox 1 gene (*Cux1*), the host gene encoding miR-721 in mice, which is annotated in the *Ensembl* genome browser^44^. Both La and LPS groups showed reduced *Cux1* expression in both strains and 4 h and 24 h (**Supplementary Figure 6**).

**Figure 3.**
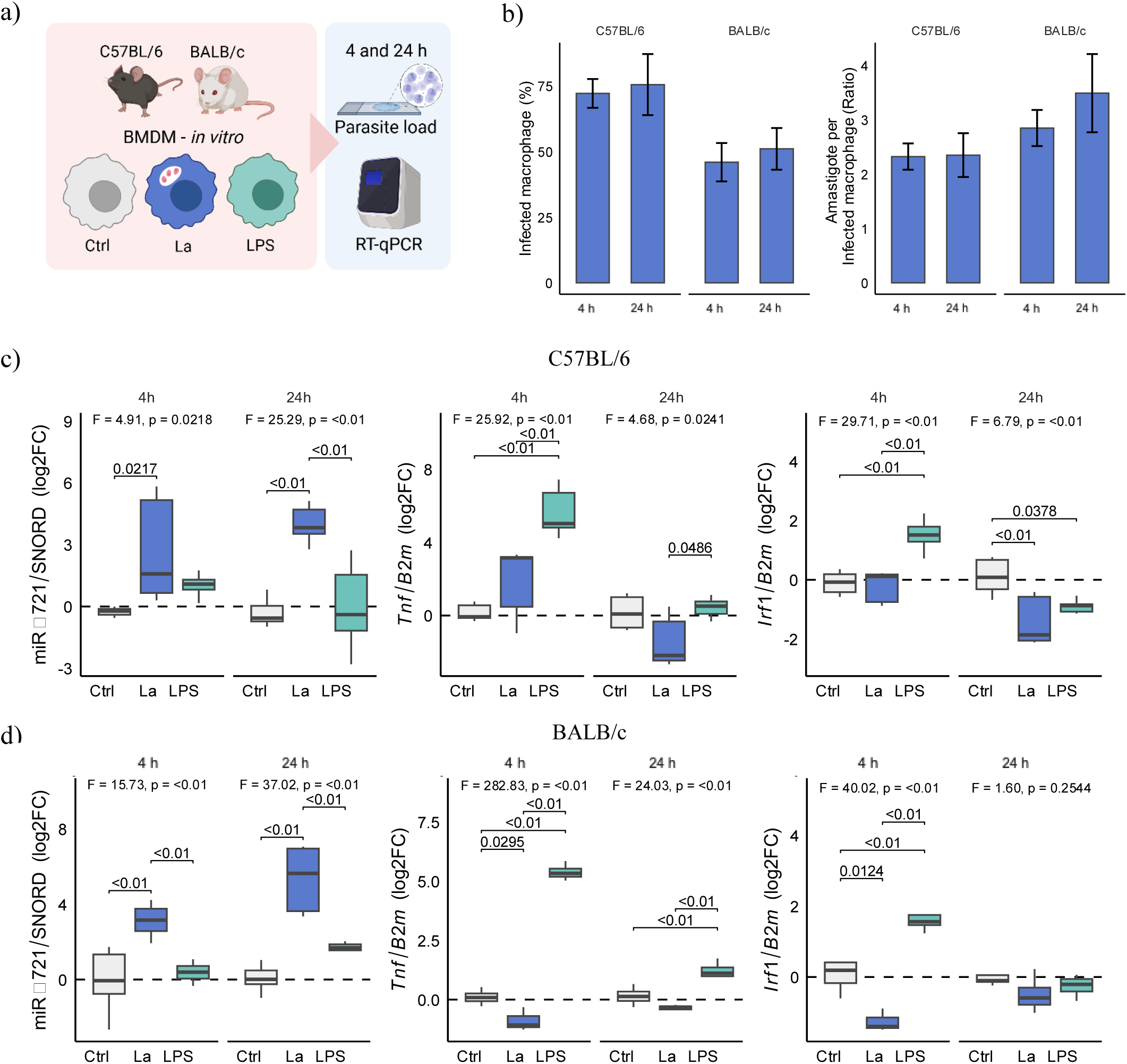
Experimental analysis of the miR-721, *Tnf* and *Irf1* expression axis in C57BL/6 and BALB/c BMDMs. **a)** Schematic overview of the validation strategy. **b**) Infection parameters showing the percentage of infected macrophages and the amastigote-to-macrophage ratio at 4 h and 24 h post infection. **c**) Expression of miR-721, *Tnf*, and *Irf1* in C57BL/6 macrophages under Ctrl, La, and LPS conditions. **d**) Expression of miR-721, *Tnf*, and *Irf1* in BALB/c macrophages under Ctrl, La, and LPS conditions.

Next, we evaluated the expression of *Tnf* and *Irf1* in BMDMs from C57BL/6 and BALB/c macrophages. In C57BL/6 macrophages, *L. amazonensis* infection did not significantly alter *Tnf* expression at 4 h compared to Ctrl conditions, whereas LPS stimulation induced *Tnf* expression at the same timepoint (**Figure 3c**). In contrast, LPS induced *Irf1* expression at 4 h, but reduced its levels at 24 h compared to Ctrl conditions. Notably, infection led to a reduction of *Irf1* expression at 24 h (**Figure 3c**). In BALB/c macrophages, La significantly reduced *Irf1* at 4h (**Figure 3d)**. On the other hand, LPS stimulation induced *Tnf* expression at 4 and 24h compared to Ctrl (**Figure 3d**).

We also analyzed *Nos2* expression, a key mediator of nitric oxide production in response to *Leishmania* infection and induced by IRF1-dependent inflammatory signaling. In C57BL/6 macrophages, LPS induced *Nos2* at 4 and 24 h compared to Ctrl and La conditions. In BALB/c macrophages, *Nos2* expression was induced exclusively by LPS at both timepoints (**Supplementary Figure S6a** and **S6b**). These findings demonstrate that *L. amazonensis* infection modulates the expression of key inflammatory genes. The observed induction of miR-721, particularly in association with the negative modulation of *Tnf* and *Irf1*, supports a potential role for miR-721 in the downregulation of inflammatory responses during infection.

### 3.3 Functional inhibition of miR-721 enhances *Tnf* and *Irf1* expression during *L. amazonensis* infection

To assess whether *Tnf* and *Irf1* are regulated by miR-721, we performed functional inhibition assays in RAW264.7 macrophages 24 h prior to infection with *L. amazonensis*. *In silico* analysis using miRmap identified conserved miR-721 binding sites within the 3′ untranslated regions (3′UTRs) of both *Tnf* (77% exact probability and miRmap score 95.30) and *Irf1* (77.56% exact probability and miRmap score 84.67), supporting their classification as putative direct miR-721 targets (**Figure 4a**). Efficient inhibition of miR-721 following macrophage transfection was confirmed by RT-qPCR, which revealed a significant reduction in miR-721 levels at 24 h post-infection compared to infected negative control (NC-miR) (**Figure 4b**).

**Figure 4.**
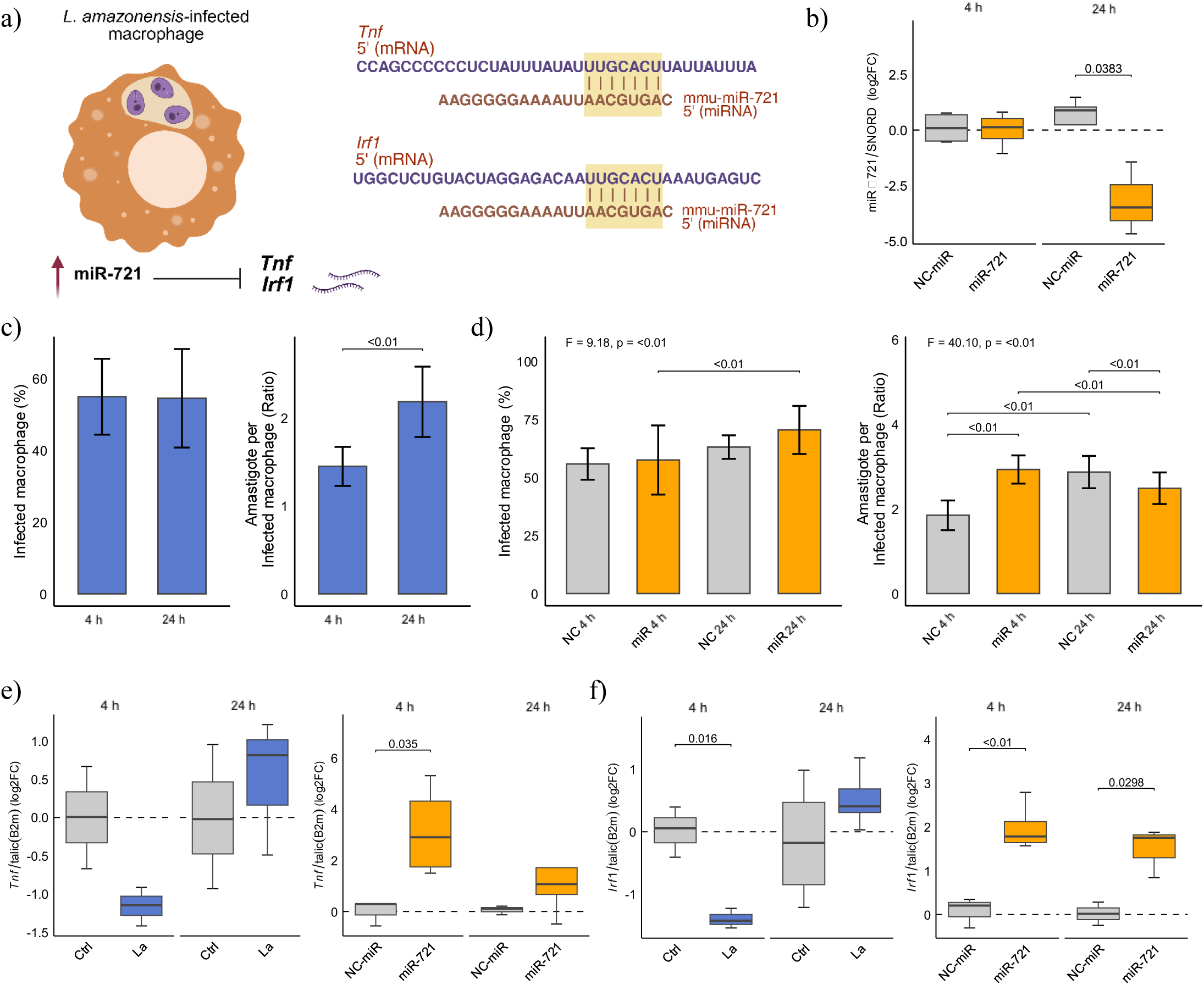
Functional validation of miR-721 inhibition and its impact on *Tnf* and *Irf1* expression. **a)** Schematic overview of the miR-721 inhibition strategy in RAW cells. **b**) Boxplot showing the expression by log2FC of miR-721 inhibition in RAW cells. **c-d)** Bar plot showing the percentage of infected RAW macrophages, amastigote-to-macrophage ratio (**c**), and miR-721 inhibition ratio (**d**) at 4 h and 24 h post infection. **e)** Box plot showing expression of *Tnf* in RAW cells under Ctrl (Gray) and La (Blue) conditions at 4 h and 24 h, and in infected RAW cells transfected with NC-miR (Gray) or miR-721 inhibitor (Orange). **f**) Box plot showing expression of *Irf1* in RAW cells under Ctrl (Gray) and La (Blue) conditions at 4 h and 24 h, and in infected RAW cells transfected with NC-miR (Gray) or miR-721 inhibitor (Orange).

Infection parameters showed around 55% of infected macrophages and at 4 and 24h and, with a marked increase in the number of amastigotes per infected macrophage from 1.5 to 2.2 at 24 h (**Figure 4c**). Importantly, miR-721 inhibition did not significantly alter infection rates during the early stages of infection, indicating that miR-721 modulation does not impair parasite entry (**Figure 4d**). Notably, the number of amastigotes per infected macrophage significantly increased at 4 h of inhibition but decreased at 24 h compared to Ctrl group (**Figure 4d**). Overall, these results suggest a dynamic effect of miR-721 inhibition on parasite burden over time (**Figure 4c** and **4d**).

In infected macrophages, *Tnf* expression showed a tendency toward downregulation at 4 h compared to the control group (**Figure 4e**). However, the miR-721 inhibition significantly increased *Tnf* expression at this timepoint (**Figure 4e**). Similarly, *Irf1* expression was reduced at 4 h post-infection compared to control, returning to baseline levels at 24 h (**Figure 4f**). This pattern is consistent with the restrained inflammatory profile observed in our previous RNAseq analysis results (**Figure 2**). Indeed, miR-721 inhibition significantly induced *Irf1* expression at 4 and 24 h post-infection compared to the Ctrl group (**Figure 4f**).

Notably, we observed reduced *Cux1* transcription at 4h in C57BL/6 macrophages and at 24h in BALB/c macrophages during infection or LPS stimulation, suggesting that miR-721 regulation may occur independently of the temporal expression of its host gene, CUX1. *Nos2* expression following infection was increased significant only at 4 h in C57BL/6 macrophages compared to Ctrl, whereas LPS stimulation induced a markedly stronger increase (**Supplementary Figure S7a** and **S7b**). Together, these experimental validation findings support a functional link between miR-721 and the regulation of pro-inflammatory response during *L. amazonensis* infection, mainly through the modulation of *Tnf* and *Irf1*.

To further explore the downstream role of TNF-IRF1 regulatory axis, we conduct a PPI analysis using the complete RNA-seq dataset without restricting the analysis to predicted miR-721 targets. This approach focused on the full DEGs set to identify genes potentially associated with TNF- and IRF1-related pathways. This strategy enabled the identification of transcriptional programs associated with macrophage immune responses during infection. Notably, the relationship between TNF and IRF1 defines a central immune axis supported by genes such as *Serpine1*, *Csf1*, *Cd69*, and *Maf*, and further included genes previously identified in the network presented in **Figure 2**, including *Gadd45a* and *Lrp8* (**Figure 5a**; **Supplementary Table S10**). Also, the TNF-IRF1 axis included genes related to autophagy, as *Atg*12 and *Atg*16*l*1, as well as apoptosis and cell survival, including *Bcl*2*l*11 and *Apaf*1. The Circos plot highlights these interactions across macrophages strains, incorporating their respective Log2FC values and showing that genes linked to TNF-IRF1 axis are more strongly upregulated in infected C57BL/6 macrophages than in BALB/c macrophages (**Figure5b**). Together, these findings support the existence of a miR-721-associated with a negative regulatory network centered on *Tnf* and *Irf1* that coordinates macrophage inflammatory responses during *L. amazonensis* infection (**Figure 6**).

**Figure 5.**
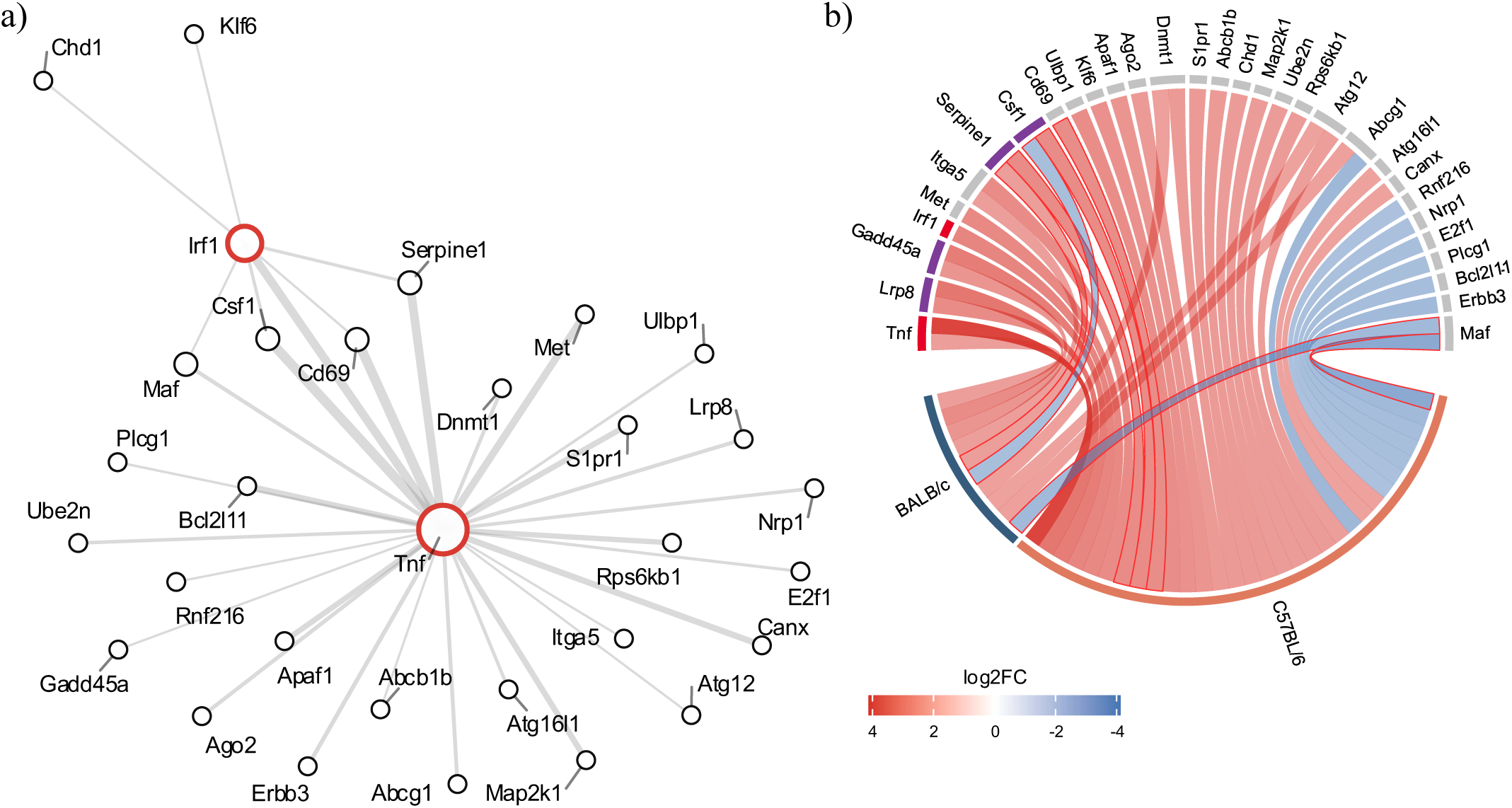
Protein-protein interaction (PPI) analysis of TNF-IRF1 interactions within the transcriptomic landscape. **a**) PPI network of DEGs from the complete transcriptomic dataset, showing interactions involving *Tnf* or *Irf1*. **b**) Circos plot representing interaction landscape. Genes interacting with both *Tnf* and *Irf1* are indicated with a red outline, while genes overlapping with the main predicted targets are marked in purple on the outer circle. *Tnf* and *Irf1* are indicated in red on the outer circle for reference.

**Figure 6.**
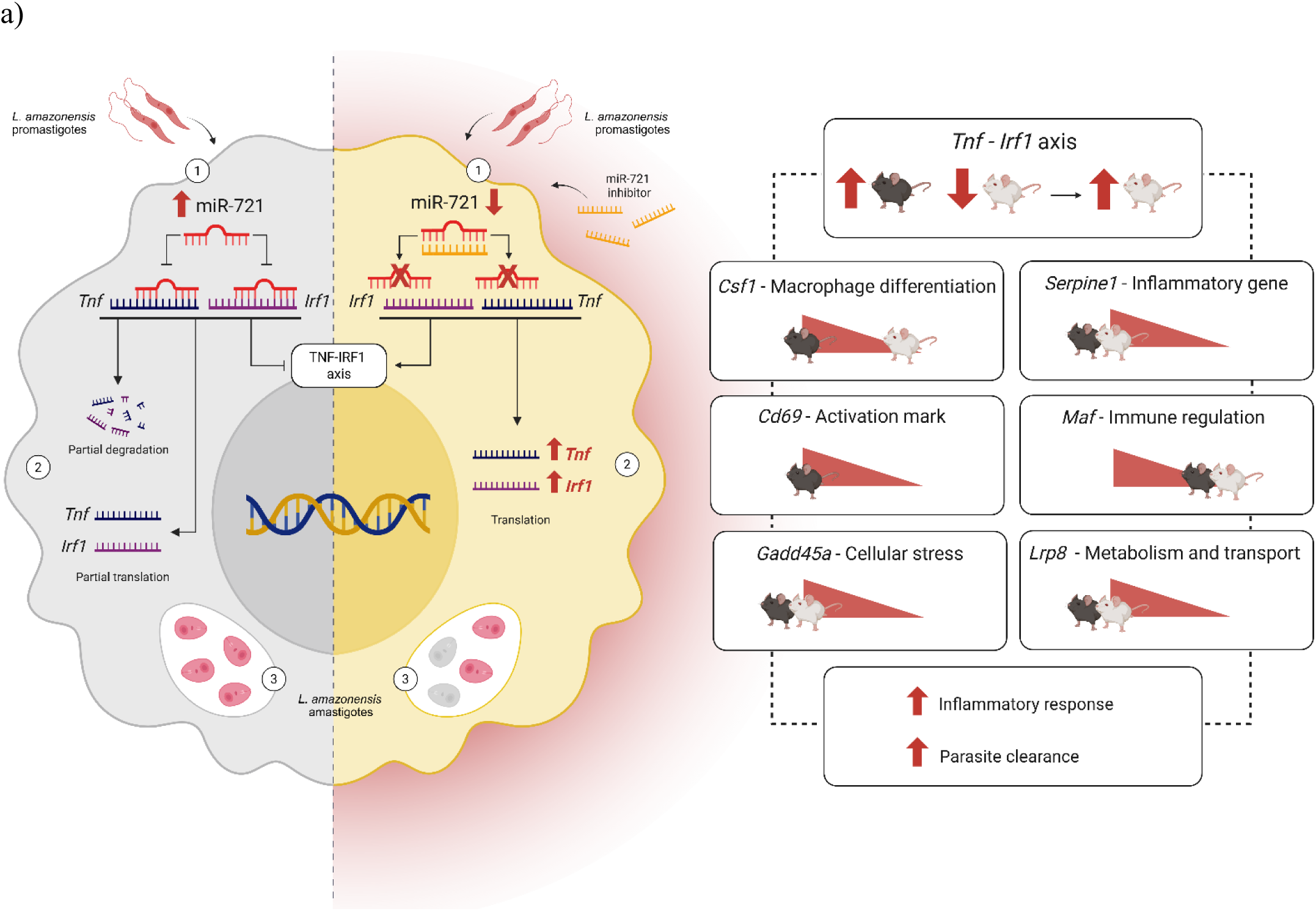
Schematic representation of miR-721–TNF–IRF1 axis. Schematic representation of the proposed regulatory interaction between miR-721 and the TNF–IRF1 axis during infection. On the left side, infection induces upregulation of miR-721, which targets *Tnf* and *Irf1* expression (1) by impairing their translation and promoting partial mRNA degradation (2), resulting in the parasite persistence within phagolysosome (3). On the right side, inhibition of miR-721 (1) restores *Tnf* and *Irf1* expression (2), enhances the inflammatory response, and promotes parasite elimination (3). miR-721 can also modulate downstream components of the TNF-IRF1 axis, such as *Serpine1, Maf, Csf1, Cd69, Lrp8* and *Gadd45a*, in C57BL/6 and BALB/c mice.

## 4 Discussion

In this work, transcriptomic correlation analysis in BMDM macrophages during *L. amazonensis* infection, combined with the screening of predicted miR-721 targets, identified *Tnf* and *Irf*1 as central regulators of the host response. The TNF-IRF1 axis may represent a central regulatory node controlling macrophage inflammatory activation and leishmanicidal responses. Our findings suggest that miR-721-mediated suppression restrains macrophage inflammatory responses, thereby favoring parasite persistence. In line with this, previous studies have shown that the knockout of *Tnf* facilitates the development of fatal cutaneous leishmaniasis infection^9,10,45–47^. During infection, parasite recognition triggers a signaling cascade via TLR2/4, MyD88, and NF-kB and IRF1^48^. IRF1 is a transcription factor responsive to IFN-γ and TNF that promotes M1-polarization^49–51^, upregulating pro-inflammatory genes, as *Caspase-1, Nos2* and *Il-12b* ^48,52–55^.

Here, we showed the upregulation of miR-721 during *L. amazonensis* infection, and its role was further supported by inhibition assays, similarly to our previous studies in *L. amazonensis*-infected macrophages^19^. In contrast to infection, LPS could not induce miR-721 expression, suggesting that miR-721 induction may occur through mechanisms that are at least partially independent of classical TLR4-mediated inflammatory activation. Our work supports a novel regulatory mechanism in *Leishmania* infection, in which miR-721 is predicted to target *Tnf* and *Irf1* to downregulate mRNA levels in infected macrophages, thereby modulating inflammatory gene expression. TNF and IRF1 can function cooperatively, with TNF providing extracellular inflammatory signaling via TNF receptor and IRF1 regulating intracellular antimicrobial transcriptional programs^56,57^. In addition, miR-721 can indirectly impact downstream components of the TNF-IRF1 axis, such as *Serpine1*, *Csf1*, *Cd69*, and *Maf* as observed in the circos plot. These genes are associated with macrophage activation, differentiation, and inflammatory regulation. For example, the downstream molecular cascade of the TNF-IRF1 axis may regulate the inflammatory activation supports the hypothesis that miR-721 promotes a macrophage profile with a tendency towards M2 over M1. Recently data shown an M2-like macrophage profile promoted via *Maf*^58,59^, chronic inflammatory lesions and cell infiltration associated with *Serpine*1^60^, cell activation observed via *Cd*69 marker, and *Csf1* induce the growth and differentiation of macrophages^61,62,58,63,64^.

Using differential gene expression analysis, we found common and specific DEGs that reflect differences between the C57BL/6 and BALB/c genotypes and the potential of *L. amazonensis* to subvert immune responses via miR-721. C57BL/6 mice are widely recognized as a relatively resistant model in *Leishmania* infection studies, displaying more robust inflammatory and antimicrobial responses ^10,45^. In agreement with this, our transcriptomic analyses showed a more organized and enriched TNF-IRF1-associated inflammatory network in C57BL/6 macrophages, whereas BALB/c macrophages exhibited a comparatively restricted inflammatory profile. In this context, the function of miR-721, combined with the set of genes identified in resistant macrophages, warrants further investigation under suppressive or chronic infection conditions.

Our enrichment analysis highlighted inflammatory-response-associated genes in both strains, which may compose a signature of this response in macrophages. The expression of *Lrp8*, *Slc25a44*, *Gadd45a, Arrdc3*, *Ampd3*, *Osbpl11*, *Maf*, and *Tnf* is upregulated in C57BL/6 and BALB/c macrophages. These genes are related to macrophage function and may trigger inflammatory responses through oxidative stress^65,66^. The disruption of glycosphingolipid biosynthesis mediated by LRP8 or lipids transport via ORP11 (encoded by Ospbl11), impairs the efficiency of uptake and antigen presentation, and protection against oxidative stress^65,67–70^. *Leishmania* can hijack these lipid rafts via lipophosphoglycans (LPGs) to avoid NADPH oxidase formation in the phagolysosome^71,72^.

The activation of the TNF-IRF-1 axis in C57BL/6 macrophages may lead to a more resistant profile, as observed in the transcriptional patterns observed in our analyses. Furthermore, the breakdown of this network in BALB/c macrophages likely facilitates the parasite persistence s. In particular, *Tnf* expression was highly upregulated in C57BL/6 macrophages but less prominently induced in BALB/c macrophages. A similar pattern occurs for *Irf1*, which was induced in C57BL/6 macrophages but absent in BALB/c macrophages

Together, these findings support the existence of a miR-721-associated regulatory network centered on *Tnf* and *Irf1* that coordinates macrophage inflammatory responses during *L. amazonensis* infection. Although our findings support a regulatory interaction between miR-721 and the TNF-IRF1 axis, additional studies using direct binding assays and *in vivo* models will be important to fully establish the mechanistic and physiological relevance of this pathway. Nonetheless, our results identify miR-721 as a previously underexplored regulator associated with macrophage inflammatory responses during *Leishmania* infection and provide new insights into mechanisms of parasite-driven immune modulation.

## Code availability

The code will be deposited in GitHub repository, which is currently being prepared and will be made publicly available upon publication.

## Data availability

The single-cell RNA sequencing data generated in this study were deposited in SRA under accession number PRJNA481042. PCR data will be available at GitHub repository on this study.

## Competing Interest Statement

The Authors declare no Competing Financial or Non-Financial Interests.

## Contributions

S.M.M. conceived the study, supervised and provided funding. J.M.Z., I.A.F.B., D.L.M.F. and S.M.M wrote the manuscript. J.M.Z. and C.A.T. designed and performed the experiments of gene expression and functional inhibition of miRNAs. I.A.F.B., E.S.A.S. and D.L.M.F. performed data curation and integration, including the development of analytical code and repository organization, RNA-seq data processing, and statistical analysis. J.M.Z., D.L.M.F., I.A.F.B., S.M.M., K.T.K., C.A.B., R.J.L. S. and M.R.I.L provided scientific assistance, intellectual support and critically revised the manuscript. J.M.Z. and I.A.F.B. contributed equally and are considered co-first authors. All authors have read and agreed to the published version of the manuscript.

Instituto Nacional de Ciência e Tecnologia em Doenças Tropicais (INCT-DT)

## Acknowledgments

We acknowledge the São Paulo Research Foundation (FAPESP grants: 2024/08802-3 and 2024/08360-0 to S.M.M.; 2025/03433-2 to D.L.M.F.; 2025/04633-5 to C.A.T.; 2022/08700-0 and 2024/06233-1 to J.M.Z.; 2025/05325-2 to I.A.F.B.; 2025/15259-7 to K.T.K.; 2022/06074-5 to E.S.A.S.), and the Coordination for the Improvement of Higher Education Personnel (CAPES) Financial Code 88887.992612/2024-00 (grant to I.A.F.B.) for financial support. We acknowledge the National Council for Scientific and Technological Development (CNPq) Brazil grant 303384/2024-7 to S.M.M and MCTI/CNPq/CAPES/FAPs Grant 408294/2024-8, Instituto Nacional de Ciência e Tecnologia em Doenças Tropicais (INCT-DT). This work has been funded by national funds, through the FAPESP grants: 2024/08802-3 and Foundation for Science and Technology (FCT) of Portugal, under projects UID/06304/2025 (https://doi.org/10.54499/UID/06304/2025) and LA/P/0050/2020 (https://doi.org/10.54499/LA/P/0050/2020).

---

**Supplementary Figure 1.**
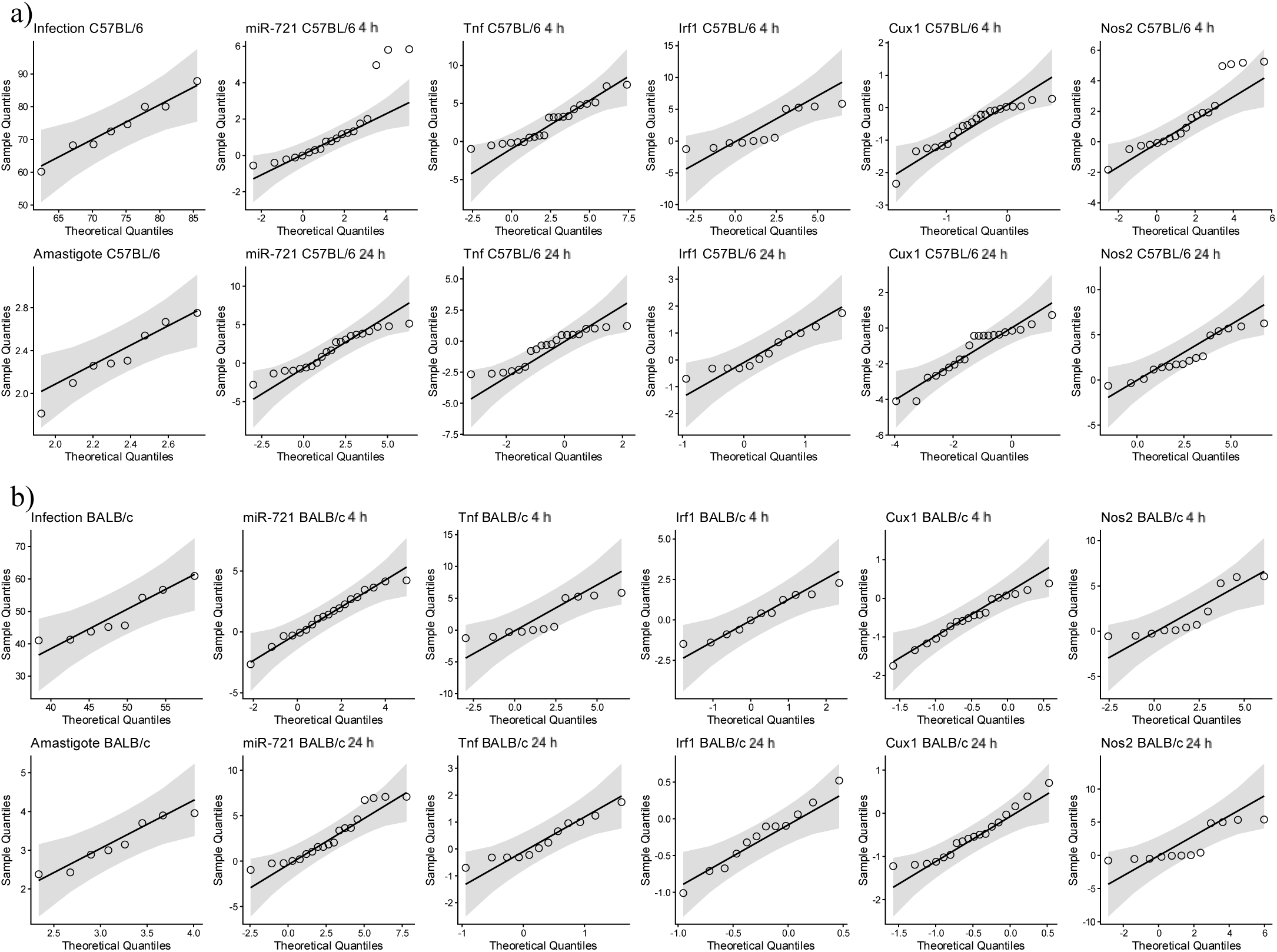
Normality assessment of infection metrics and gene expression in C57BL/6 and BALB/c BMDMs. **a**) QQ plots showing the distribution of values for infection percentage, amastigote rate, miR-721, *Tnf*, *Irf1*, *Cux1*, and *Nos2* in C57BL/6 BMDMs at 4 h and 24 h. **b**) QQ plots showing the distribution of values for infection percentage, amastigote rate, miR-721, *Tnf*, *Irf1*, *Cux1*, and *Nos2* in BALB/c BMDMs at 4 h and 24 h.

**Supplementary Figure 2.**
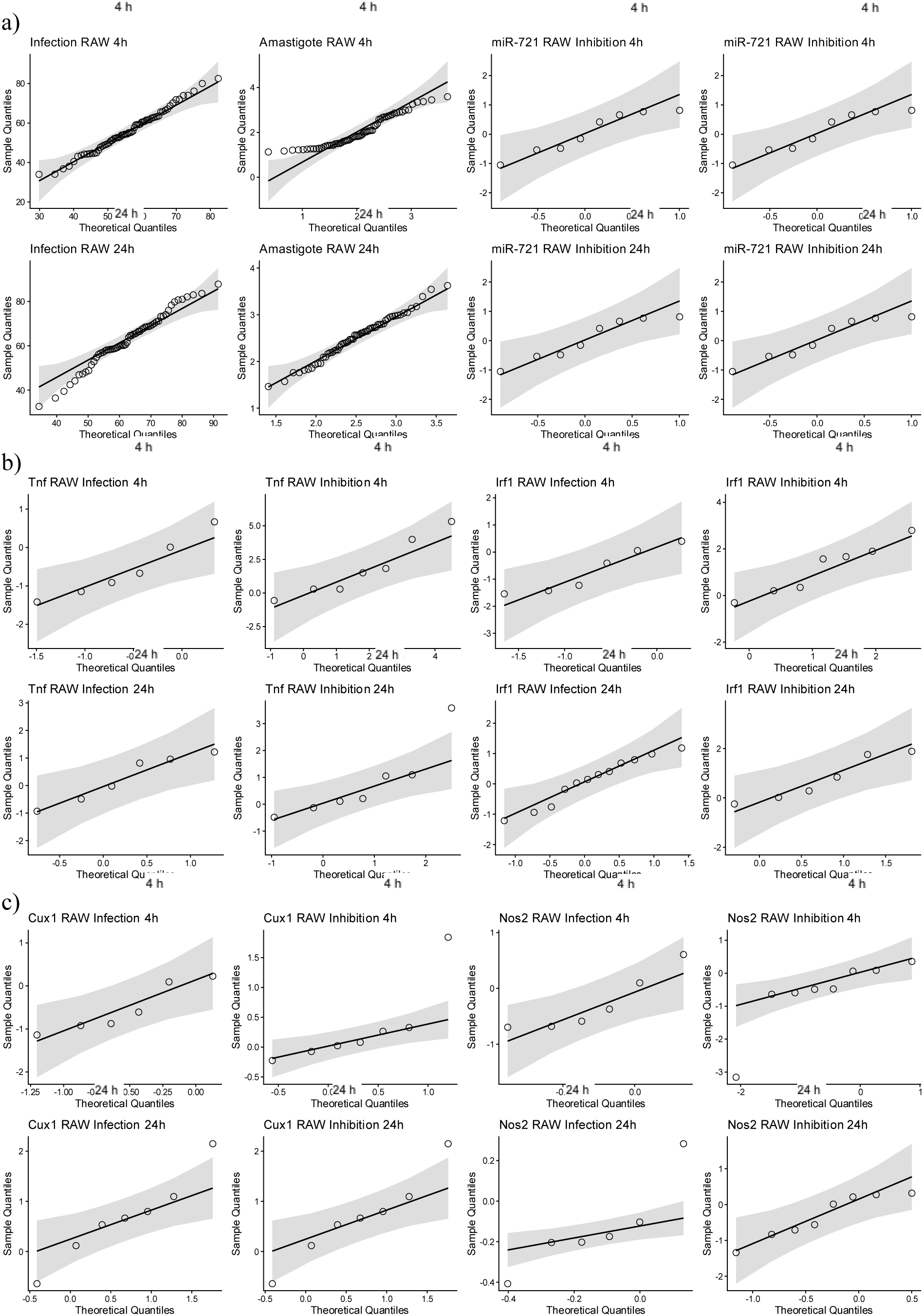
Normality assessment of infection metrics and gene expression in RAW cells. **a**) QQ plots showing the distribution of values for infection percentage, amastigote rate, miR-721, *Tnf*, *Irf1*, *Cux1*, and *Nos2* expression in RAW cells at 4 h and 24 h, in infection and inhibition condition. **b**) QQ plots showing the distribution of values for *Tnf* and *Irf1* expression in RAW cells at 4 h and 24 h, in infection and inhibition condition. **c)** QQ plots showing the distribution of values for *Cux1* and *Nos2* expression in RAW cells at 4 h and 24 h, in infection and inhibition condition.

**Supplementary Figure 3.**
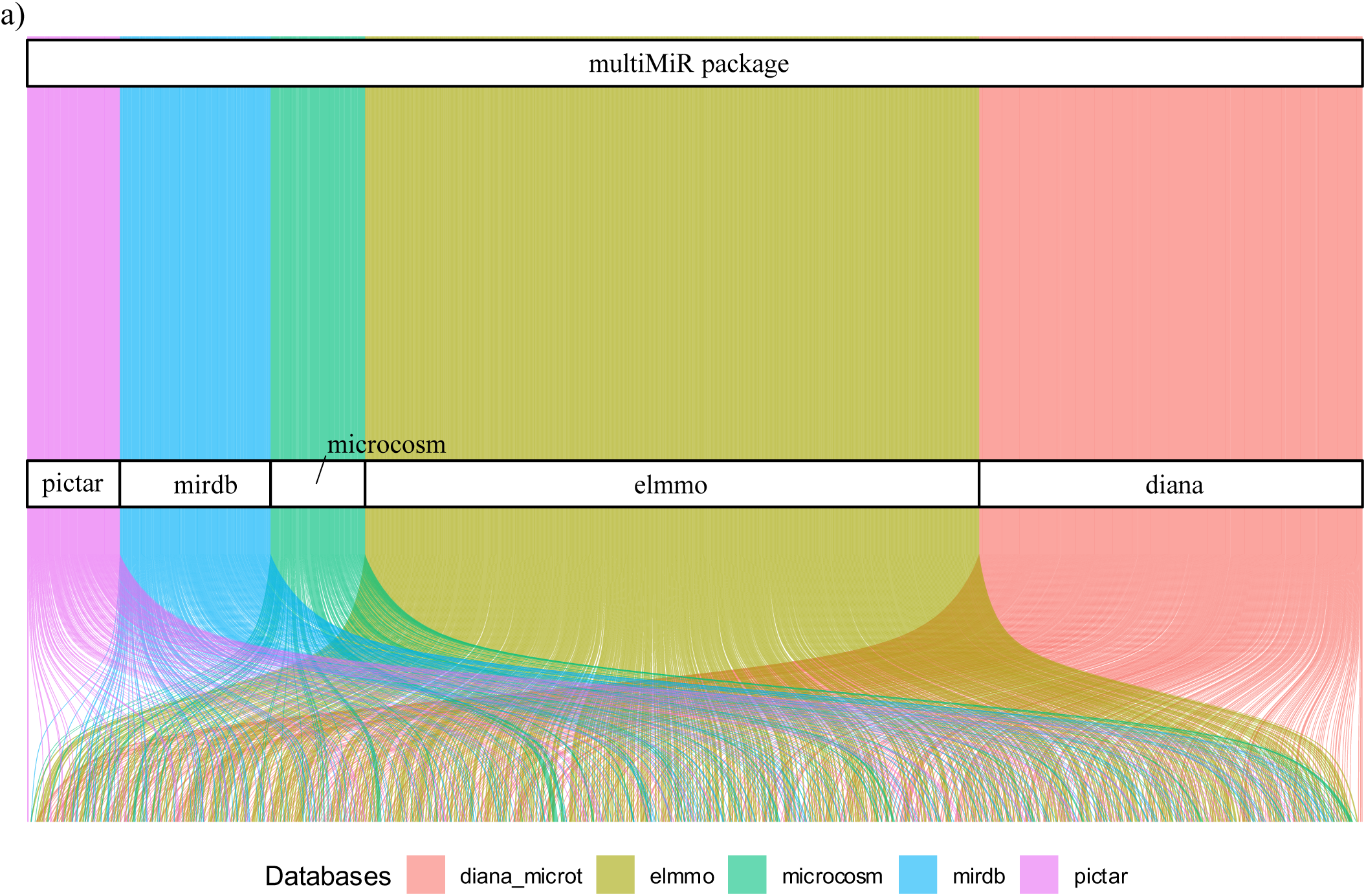
Mapping of miR-721 predicted targets. All predicted targets identified for miR-721 are shown according to database of origin and intersection patterns. The alluvial illustrates the proportion of genes supported by multiple databases, highlighting high-confidence targets that were subsequently used to filter RNA-seq dataset.

**Supplementary Figure 4.**
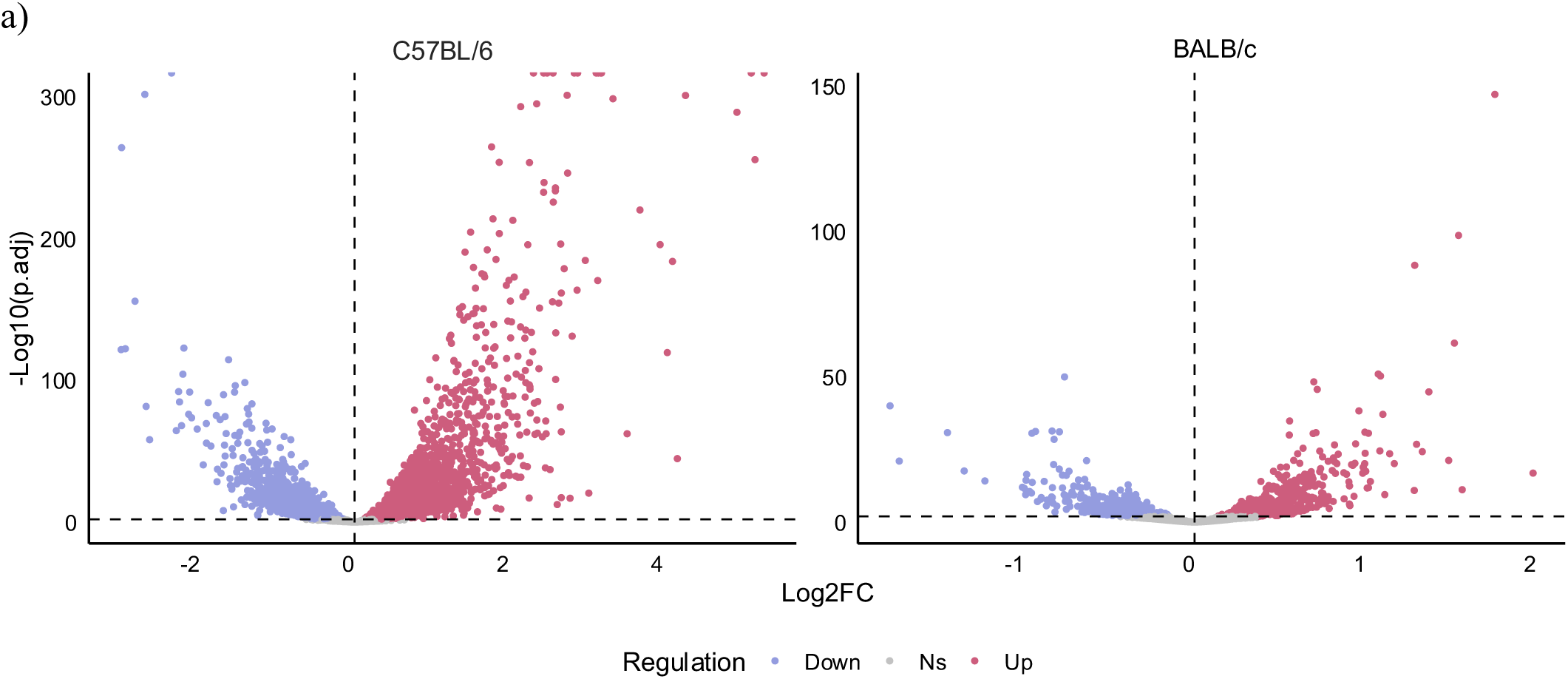
Differential expression profile of the complete bulk RNAseq. Volcano plots for C57BL/6 and BALB/c infected macrophages showing global transcriptional responses to *L. amazonensis*. These plots provide the unfiltered landscape, illustrating the broader gene modulation from which miR-721 targets represent a biologically meaningful subset.

**Supplementary Figure 5.**
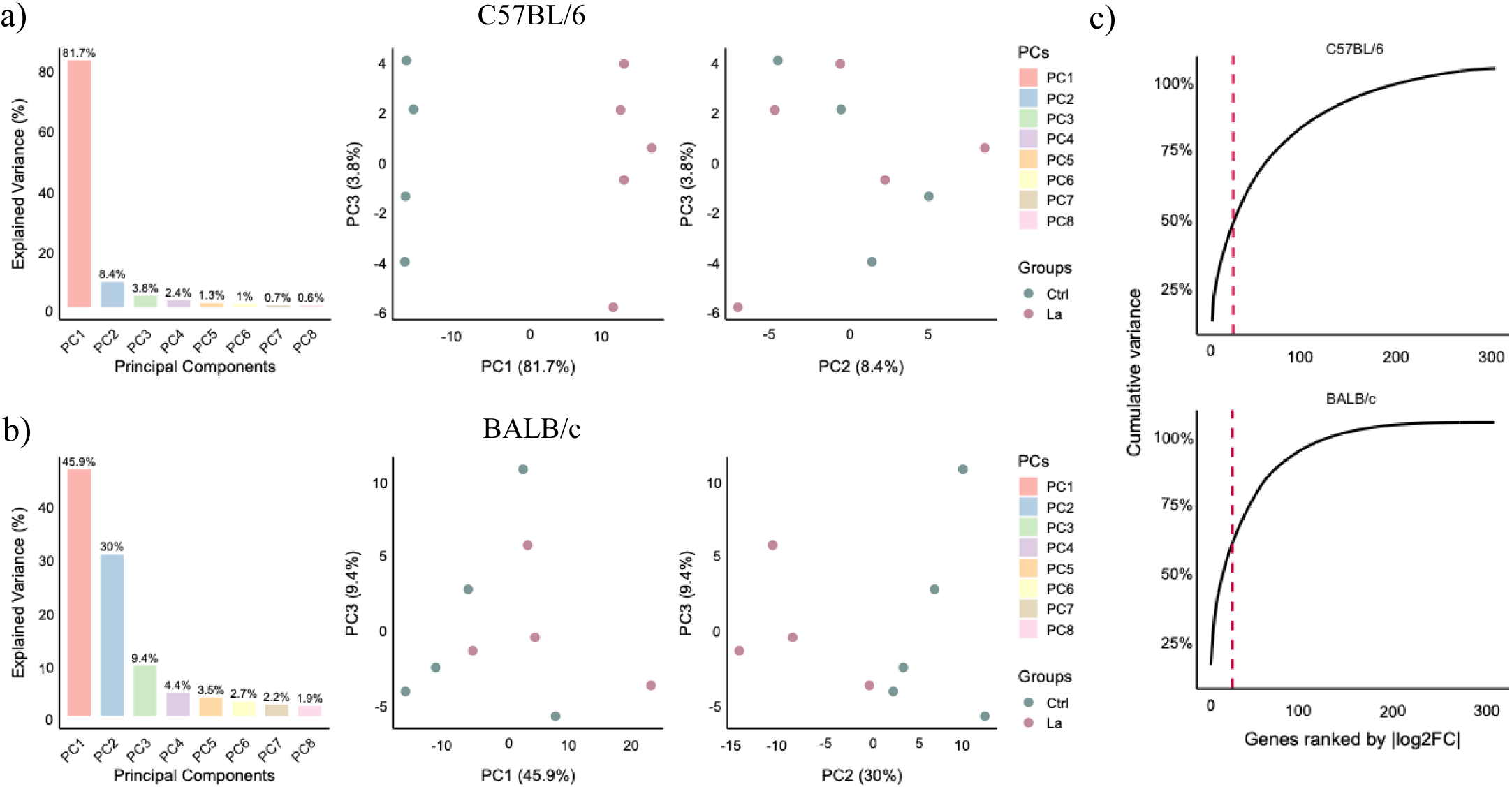
PCA deepening and gene Log2FC cumulative variance filtering. **a–b)** Barplots showing the percentage of variance explained by each principal component for the C57BL/6 and BALB/c datasets. The elbow rule was applied to guide PC selection, resulting in the inclusion of components up to PC3 for both strains. Scatter plots illustrate the corresponding PCA projections, displaying all dimensions required to capture the major sources of variation between groups. **c)** Cumulative variance plot illustrating the stepwise selection of genes contributing up to 90% of Log2FC cumulative variance, forming the final set used for enrichment and network analyses.

**Supplementary Figure 6.**
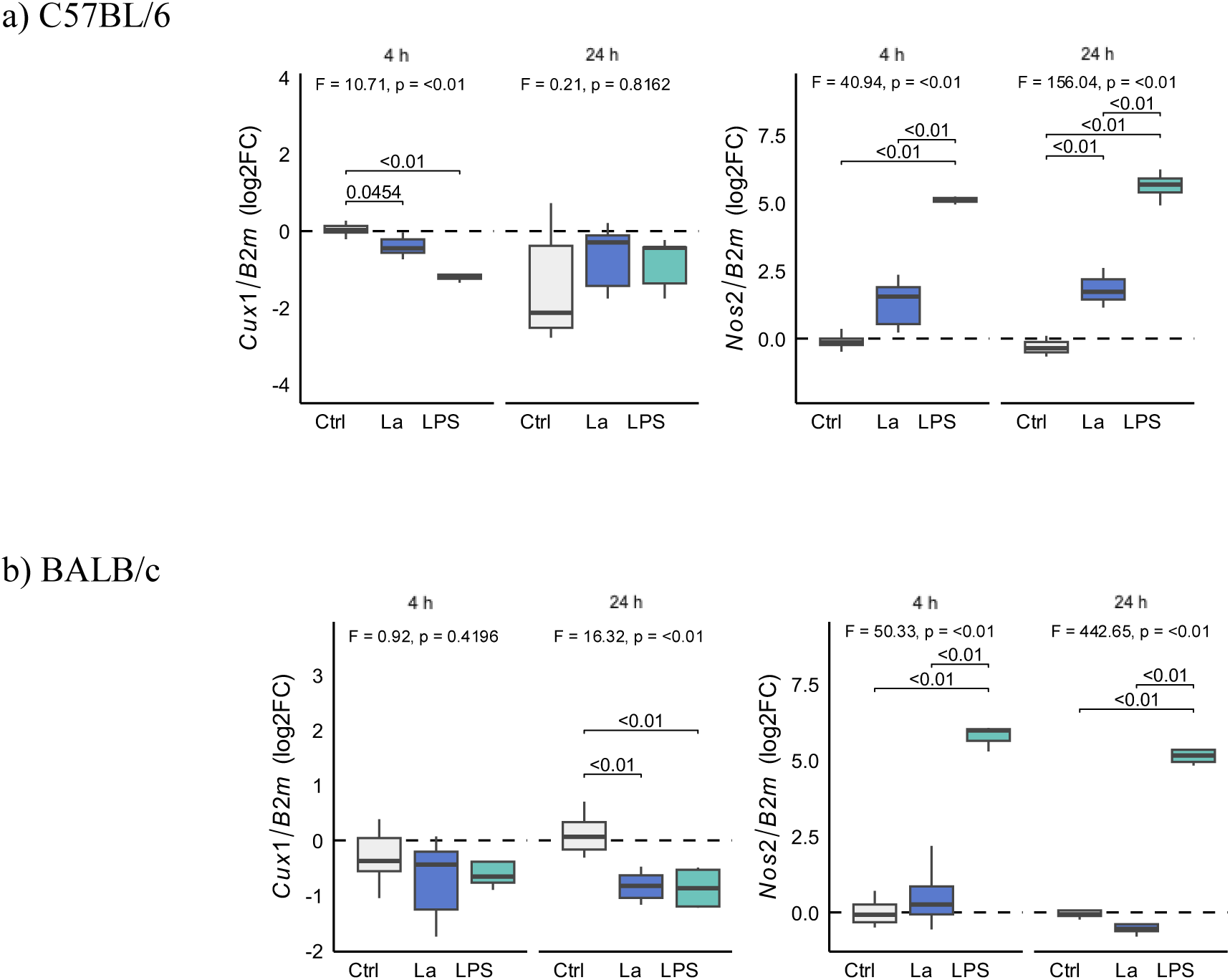
Experimental validation of Cux1 and Nos2 expression in C57BL/6 and BALB/c BMDMs. **a**) Relative expression of *Cux1* and *Nos2* in C57BL/6 macrophages under Ctrl, La, and LPS conditions. **b**) Relative expression of *Cux1* and *Nos2* in BALB/c macrophages under Ctrl, La, and LPS conditions.

**Supplementary Figure 7.**
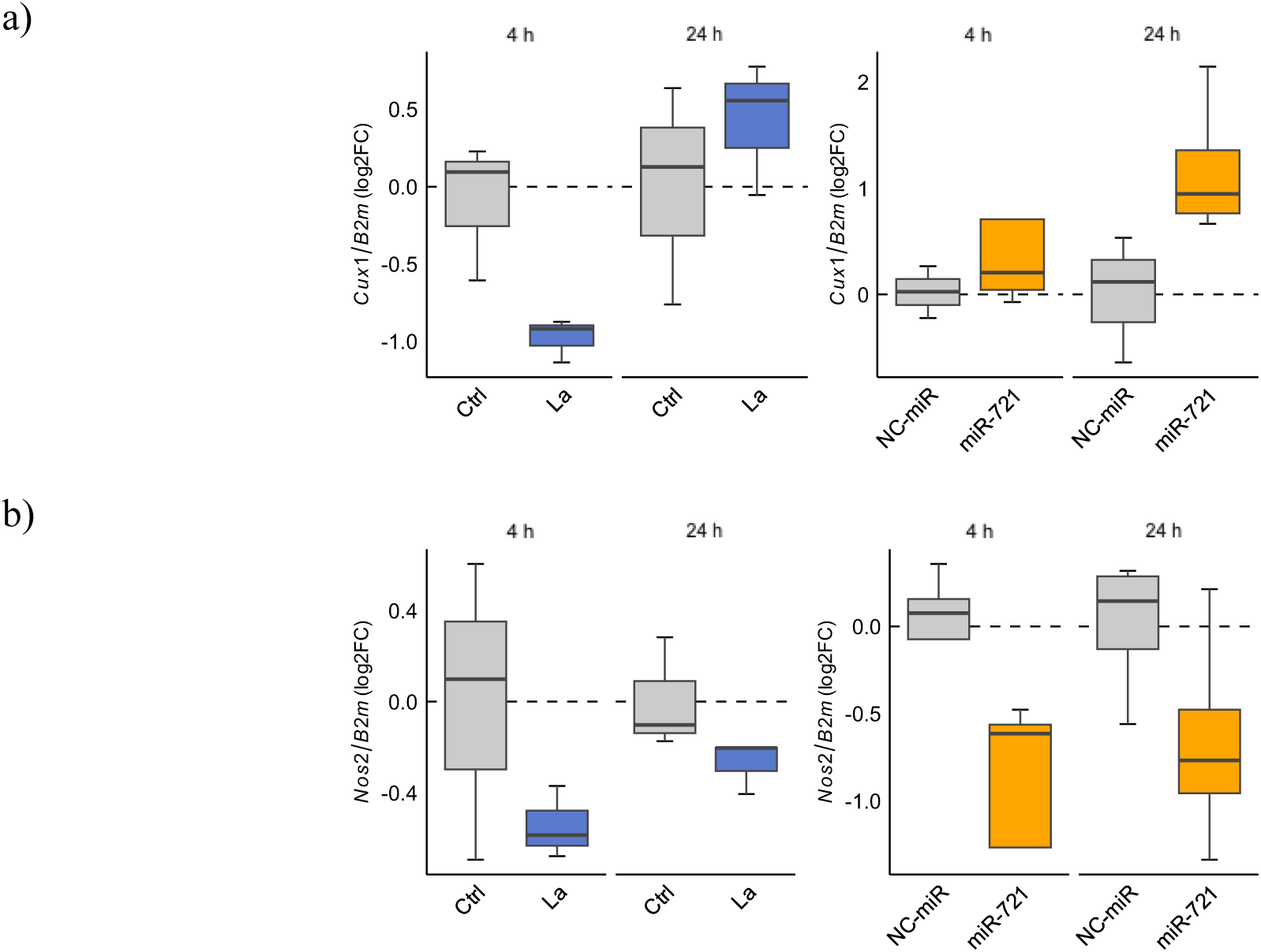
Functional validation of miR-721 inhibition and its impact on *Cux1* and *Nos2* expression. **a**) Relative expression of *Cux1* in RAW cells under Ctrl and La conditions at 4 h and 24 h, and in infected RAW cells transfected with NC-miR or miR-721 inhibitor. **b**) Relative expression of *Nos2* in RAW cells under Ctrl and La conditions at 4 h and 24 h, and in infected RAW cells transfected with NC-miR or miR-721 inhibitor.

## Notes

### Competing Interest Statement

The authors have declared no competing interest.

## References

1. Burza, S., Croft, S. L. & Boelaert, M. Leishmaniasis. The Lancet 392, 951–970 (2018).

2. Alvar, J. et al. Leishmaniasis Worldwide and Global Estimates of Its Incidence. PLoS ONE 7, e35671–e35671 (2012).

3. Liese, J., Schleicher, U. & Bogdan, C. The innate immune response against Leishmania parasites. Immunobiology 213, 377–387 (2008).

4. Von Stebut, E. et al. Interleukin 1α promotes TH1 differentiation and inhibits disease progression in Leishmania major-susceptible BALB/c mice. J. Exp. Med. 198, 191–199 (2003).

5. Kropf, P. et al. Toll-like receptor 4 contributes to efficient control of infection with the protozoan parasite Leishmania major. Infect Immun 72, 1920–1928 (2004).

6. Lohoff, M. et al. Interferon Regulatory Factor-1 Is Required for a T Helper 1 Immune Response In Vivo. Immunity 6, 681–689 (1997).

7. Bogdan, C. Mechanisms and consequences of persistence of intracellular pathogens: leishmaniasis as an example. Cell Microbiol 10, 1221–1234 (2008).

8. Resende, M. et al. Leishmania -Infected MHC Class II high Dendritic Cells Polarize CD4 + T Cells toward a Nonprotective T-bet + IFN-γ + IL-10 + Phenotype. J. Immunol. 191, 262–273 (2013).

9. Belkaid, Y. et al. The Role of Interleukin (IL)-10 in the Persistence of Leishmania major in the Skin after Healing and the Therapeutic Potential of Anti–IL-10 Receptor Antibody for Sterile Cure. J. Exp. Med. 194, 1497–1506 (2001).

10. Pérez-Cabezas, B. et al. Understanding Resistance vs. Susceptibility in Visceral Leishmaniasis Using Mouse Models of Leishmania infantum Infection. Front. Cell. Infect. Microbiol. 9, 30–30 (2019).

11. Palacios, G. et al. Gene Expression Profiling of Classically Activated Macrophages in Leishmania infantum Infection: Response to Metabolic Pre-Stimulus with Itaconic Acid. Trop. Med. Infect. Dis. 8, 264 (2023).

12. Bichiou, H., Bouabid, C., Rabhi, I. & Guizani-Tabbane, L. Transcription Factors Interplay Orchestrates the Immune-Metabolic Response of Leishmania Infected Macrophages. Front. Cell. Infect. Microbiol. 11, 660415 (2021).

13. Arens, K. et al. Anti-Tumor Necrosis Factor α Therapeutics Differentially Affect Leishmania Infection of Human Macrophages. Front. Immunol. 9, (2018).

14. Platanias, L. C. Mechanisms of type-I- and type-II-interferon-mediated signalling. Nat. Rev. Immunol. 5, 375–386 (2005).

15. Wang, L. et al. The multiple roles of interferon regulatory factor family in health and disease. Signal Transduct. Target. Ther. 9, 282 (2024).

16. Lecoeur, H. et al. Targeting Macrophage Histone H3 Modification as a Leishmania Strategy to Dampen the NF-κB/NLRP3-Mediated Inflammatory Response. Cell Rep. 30, 1870–1882.e4 (2020).

17. Manzano-Román, R. & Siles-Lucas, M. MicroRNAs in parasitic diseases: Potential for diagnosis and targeting. Mol. Biochem. Parasitol. 186, 81–86 (2012).

18. Acuna, S. M., Floeter-Winter, L. M. & Muxel, S. M. MicroRNAs: Biological Regulators in Pathogen-Host Interactions. Cells 9, (2020).

19. Muxel, S. M., Laranjeira-Silva, M. F., Zampieri, R. A. & Floeter-Winter, L. M. Leishmania (Leishmania) amazonensis induces macrophage miR-294 and miR-721 expression and modulates infection by targeting NOS2 and L-arginine metabolism. Sci. Rep. 7, 44141 (2017).

20. Kabeer, S. W., Sharma, S., Sriramdasu, S. & Tikoo, K. MicroRNA-721 regulates gluconeogenesis via KDM2A-mediated epigenetic modulation in diet-induced insulin resistance in C57BL/6J mice. Biol. Res. 57, 27–27 (2024).

21. Zhang, J., Wang, N. & Xu, A. Cmah deficiency may lead to age-related hearing loss by influencing miRNA-PPAR mediated signaling pathway. PeerJ 7, e6856–e6856 (2019).

22. Feiz Haddad Hossein, Nourian, A. H. H. M. H. R. Expression of MicroRNA of Macrophages Infected with Attenuated Leishmania major Parasite. J. Pediatr. Infect. Dis. 16, 106–110 (2021).

23. Ke, B. et al. Astragalus polysaccharides attenuates TNF-α-induced insulin resistance via suppression of miR-721 and activation of PPAR-γ and PI3K/AKT in 3T3-L1 adipocytes. Am. J. Transl. Res. 9, 2195–2206 (2017).

24. Aoki, J. I. I. et al. Differential immune response modulation in early Leishmania amazonensis infection of BALB/c and C57BL/6 macrophages based on transcriptome profiles. Sci Rep 9, 19841 (2019).

25. Love, M. I., Huber, W. & Anders, S. Moderated estimation of fold change and dispersion for RNA-seq data with DESeq2. Genome Biol. 15, 550 (2014).

26. Love, M., Ahlmann-Eltze, C., Forbes, K., Anders, S. & Huber, W. DESeq2: differential gene expression analysis package. Bioconductor. *Bioconductor* http://bioconductor.org/packages/DESeq2/.

27. Wickham, H. Ggplot2: Elegant Graphics for Data Analysis. (Springer International Publishing, New York, NY, 2016).

28. R: The R Project for Statistical Computing. https://www.r-project.org/.

29. Ru, Y. et al. The multiMiR R package and database: integration of microRNA–target interactions along with their disease and drug associations. Nucleic Acids Res. 42, e133 (2014).

30. Robinson, M. D., McCarthy, D. J. & Smyth, G. K. edgeR: a Bioconductor package for differential expression analysis of digital gene expression data. Bioinformatics 26, 139–140 (2010).

31. Das, S. & Rai, S. N. Statistical Approach for Biologically Relevant Gene Selection from High-Throughput Gene Expression Data. Entropy 22, 1205 (2020).

32. Yu, G. Thirteen years of clusterProfiler. The Innovation 5, 100722 (2024).

33. Xu, S. et al. Using clusterProfiler to characterize multiomics data. Nat. Protoc. 19, 3292–3320 (2024).

34. Wu, T. et al. clusterProfiler 4.0: A universal enrichment tool for interpreting omics data. The Innovation 2, 100141 (2021).

35. Yu, G., Wang, L.-G., Han, Y. & He, Q.-Y. clusterProfiler: an R Package for Comparing Biological Themes Among Gene Clusters. OMICS J. Integr. Biol. 16, 284–287 (2012).

36. Szklarczyk, D. et al. The STRING database in 2023: protein–protein association networks and functional enrichment analyses for any sequenced genome of interest. Nucleic Acids Res. 51, D638–D646 (2023).

37. Szklarczyk, D. et al. The STRING database in 2025: protein networks with directionality of regulation. Nucleic Acids Res. 53, D730–D737 (2025).

38. Muxel, S. M., Acuña, S. M., Aoki, J. I., Zampieri, R. A. & Floeter-Winter, L. M. Toll-Like Receptor and miRNA-let-7e Expression Alter the Inflammatory Response in Leishmania amazonensis-Infected Macrophages. Front. Immunol. 9, (2018).

39. Acuña, S. M., Zanatta, J. M., de Almeida Bento, C., Floeter-Winter, L. M. & Muxel, S. M. miR-294 and miR-410 Negatively Regulate Tnfa, Arginine Transporter Cat1/2, and Nos2 mRNAs in Murine Macrophages Infected with Leishmania amazonensis. Non-Coding RNA 8, 17–17 (2022).

40. Zanatta, J. M. et al. Putrescine supplementation shifts macrophage L-arginine metabolism related-genes reducing Leishmania amazonensis infection. PLOS ONE 18, e0283696–e0283696 (2023).

41. Acuña, S. M. et al. Arginase expression modulates nitric oxide production in Leishmania (Leishmania) amazonensis. PLOS ONE 12, e0187186 (2017).

42. Vejnar, C. E. & Zdobnov, E. M. MiRmap: comprehensive prediction of microRNA target repression strength. Nucleic Acids Res. 40, 11673–11683 (2012).

43. R Core Team. (2025) R: A Language and Environment for Statistical Computing. R Foundation for Statistical Computing. https://www.r-project.org/.

44. Gene: Cux1 (ENSMUSG00000029705) - Summary - Mus_musculus - Ensembl genome browser 115. http://www.ensembl.org/Mus_musculus/Gene/Summary?g=ENSMUSG00000029705;r=5:136276989-136596344.

45. Sacks, D. & Noben-Trauth, N. The immunology of susceptibility and resistance to Leishmania major in mice. Nat. Rev. Immunol. 2, 845–858 (2002).

46. Aoki, J. I. et al. Dual transcriptome analysis reveals differential gene expression modulation influenced by leishmania arginase and host genetic background. *Microb*. Genomics 6, 1–13 (2020).

47. Wilhelm, P. et al. Rapidly Fatal Leishmaniasis in Resistant C57BL/6 Mice Lacking TNF. J. Immunol. 166, 4012–4019 (2001).

48. Song, R. et al. IRF1 governs the differential interferon-stimulated gene responses in human monocytes and macrophages by regulating chromatin accessibility. Cell Rep. 34, 108891–108891 (2021).

49. Kollias, G. & Sfikakis, P. P. TNF Pathophysiology: Molecular and Cellular Mechanisms*, Vol.* 11. (Karger Medical and Scientific Publishers, 2010).

50. Bonelli, M. et al. IRF1 is critical for the TNF-driven interferon response in rheumatoid fibroblast-like synoviocytes. Exp. Mol. Med. 51, 1–11 (2019).

51. Vila-del Sol, V., Punzón, C. & Fresno, M. IFN-γ-Induced TNF-α Expression Is Regulated by Interferon Regulatory Factors 1 and 8 in Mouse Macrophages1. J. Immunol. 181, 4461–4470 (2008).

52. Chu, Y. et al. Irf1- and Egr1-activated transcription plays a key role in macrophage polarization: A multiomics sequencing study with partial validation. Int. Immunopharmacol. 99, 108072 (2021).

53. Kamijo, R. et al. Requirement for Transcription Factor IRF-1 in NO Synthase Induction in Macrophages. Science 263, 1612–1615 (1994).

54. Gao, J. et al. IRF-1 transcriptionally upregulates PUMA, which mediates the mitochondrial apoptotic pathway in IRF-1-induced apoptosis in cancer cells. Cell Death Differ. 17, 699–709 (2010).

55. Bowie, M. L. et al. Interferon-regulatory factor-1 is critical for tamoxifen-mediated apoptosis in human mammary epithelial cells. Oncogene 23, 8743–8755 (2004).

56. Yarilina, A., Park-Min, K.-H., Antoniv, T., Hu, X. & Ivashkiv, L. B. TNF activates an IRF1-dependent autocrine loop leading to sustained expression of chemokines and STAT1-dependent type I interferon-response genes. Nat. Immunol. 9, 378–387 (2008).

57. Salim, T., Sershen, C. L. & May, E. E. Investigating the Role of TNF-α and IFN-γ Activation on the Dynamics of iNOS Gene Expression in LPS Stimulated Macrophages. PLOS ONE 11, e0153289 (2016).

58. Liu, M. et al. Transcription factor c-Maf is a checkpoint that programs macrophages in lung cancer. J. Clin. Invest. 130, 2081–2096.

59. Kang, K. et al. Interferon-γ Represses M2 Gene Expression in Human Macrophages by Disassembling Enhancers Bound by the Transcription Factor MAF. Immunity 47, 235–250.e4 (2017).

60. Ju, Y. et al. Pan-cancer analysis of SERPINE1 with a concentration on immune therapeutic and prognostic in gastric cancer. J. Cell. Mol. Med. 28, e18579 (2024).

61. Ying, W., Cheruku, P. S., Bazer, F. W., Safe, S. H. & Zhou, B. Investigation of Macrophage Polarization Using Bone Marrow Derived Macrophages. J. Vis. Exp. JoVE 50323 (2013).

62. Tomiotto-Pellissier, F. et al. Macrophage Polarization in Leishmaniasis: Broadening Horizons. Front. Immunol. 9, 2529 (2018).

63. Kim, H. The transcription factor MafB promotes anti-inflammatory M2 polarization and cholesterol efflux in macrophages. Sci. Rep. 7, 7591 (2017).

64. Huang, X. et al. Immune-Related Gene SERPINE1 Is a Novel Biomarker for Diffuse Lower-Grade Gliomas via Large-Scale Analysis. Front. Oncol. 11, 646060 (2021).

65. Majumder, S. et al. Leishmania-Induced Biphasic Ceramide Generation in Macrophages Is Crucial for Uptake and Survival of the Parasite. J. Infect. Dis. 205, 1607–1616 (2012).

66. Riitano, G. et al. Role of Lipid Rafts on LRP8 Signaling Triggered by Anti-β2-GPI Antibodies in Endothelial Cells. Biomedicines 11, 3135–3135 (2023).

67. Morrison, T. A. et al. Selective requirement of glycosphingolipid synthesis for natural killer and cytotoxic T cells. Cell 188, 3497–3512.e16-3497-3512.e16 (2025).

68. Dei Cas, M., et al. Convenient and Sensitive Measurement of Lactosylceramide Synthase Activity Using Deuterated Glucosylceramide and Mass Spectrometry. Int. J. Mol. Sci. 24, 5291–5291 (2023).

69. Waltmann, M. D., Basford, J. E., Konaniah, E. S., Weintraub, N. L. & Hui, D. Y. Apolipoprotein E receptor-2 deficiency enhances macrophage susceptibility to lipid accumulation and cell death to augment atherosclerotic plaque progression and necrosis. Biochim. Biophys. Acta BBA - Mol. Basis Dis. 1842, 1395–1405 (2014).

70. Sadras, V., Petri, M. A., Jones, S. R., Peterlin, B. L. & Chatterjee, S. Glycosphingolipid-associated β-1,4 galactosyltransferase is elevated in patients with systemic lupus erythematosus. Lupus Sci. Med. 7, e000368–e000368 (2020).

71. Winberg, M. E., et al. *Leishmania donovani* lipophosphoglycan inhibits phagosomal maturation via action on membrane rafts. Microbes Infect. 11, 215–222 (2009).

72. Lodge, R., Diallo, T. O. & Descoteaux, A. Leishmania donovani lipophosphoglycan blocks NADPH oxidase assembly at the phagosome membrane. Cell. Microbiol. 8, 1922–1931 (2006).

